# Predictability and controllability shape aversive learning and stress responses through independent computational mechanisms

**DOI:** 10.64898/2026.08.26.745064

**Authors:** Daniyal Rajput, Kim Felmingham, Chin-Hsuan Sophie Lin, Marta I. Garrido

## Abstract

**BACKGROUND:** An individual’s adaptation to threatening environments under uncertainty is reflected in stress responses. Predictability (the ability to anticipate events) and controllability (the ability to control outcomes) are central to how one adapts, yet their joint influence on aversive learning remains unclear.

**METHODS:** Thirty healthy adults completed a probabilistic aversive learning task in which cue–outcome contingencies varied across levels of predictability and controllability, i.e. whether shock intensity depended on prediction accuracy. Prediction accuracy, reaction time, subjective stress ratings, and skin conductance responses were recorded throughout. Trial-wise learning dynamics were estimated using the Volatile Kalman Filter.

**RESULTS:** Prediction accuracy reduced as environments became less predictable and negatively associated with higher learning rates across predictability levels, with the strongest relationship observed in highly predictable blocks. Skin conductance responses showed that moderately predictable environments elicited responses like those in highly predictable environments when accurate predictions reduced shock intensity, but resembled responses in unpredictable environments when shock intensity was uncontrollable. Model comparison revealed a double dissociation between subjective stress ratings and skin conductance responses. Subjective ratings were best explained by model-derived volatility when prediction accuracy determined shock intensity and by belief uncertainty when it was independent of prediction accuracy, whereas skin conductance responses showed the reverse pattern. Reaction times were best explained by belief uncertainty when predictions influenced shock intensity. Higher anxiety was associated with elevated learning rates in highly and moderately predictable blocks when predictions did not control shock intensity.

**CONCLUSIONS:** These findings indicate that predictability and controllability shape aversive learning and stress responses through computational mechanisms.

## Introduction

Stress is a coordinated psychophysiological response that enables organisms to adapt to environmental challenge [1, 2], yet the mechanisms through which environmental structure shapes stress responses remain unclear. Recent work suggests that stress reflects ongoing computational inferences about environmental structure, linking behavioural, subjective, and physiological responses to internal estimates that evolve over time [3–6]. Predictability and controllability are key properties of the environment that influence inference and stress. Predictability refers to whether future outcomes can be anticipated from available cues [7], whereas controllability refers to whether outcomes depend on the individual’s actions [8]. Although both dimensions are known to shape stress responses, they have largely been studied separately [6, 9]. This is a critical limitation because real-world environments are defined by their joint structure. Understanding stress therefore requires a framework that explains how controllability and predictability are integrated to shape behaviour and physiology.

Controllability determines whether individuals can act on the environment to regulate outcomes. When control is available, stress responses are typically reduced, accompanied by greater behavioural flexibility and engagement of prefrontal systems that regulate threat processing [10–12]. In contrast, uncontrollable stress is associated with heightened anxiety, impaired learning, and behavioural passivity [13–16]. Predictability shapes how threat unfolds over time. Predictable stressors allow anticipatory preparation and temporally constrained responding, whereas unpredictable environments sustain vigilance and prolong autonomic activation [17–19]. Importantly, predictability and controllability are distinct properties of the environment, but their effects on stress responses can both differ from, and interact with, one another. A realistic mechanistic account of stress must therefore explain how these dimensions jointly shape behavioural and physiological responses.

A hierarchical Bayesian model of stress [20], which formalises how people learn to mitigate stress-inducing challenges by inferring environmental structure from experience, can provide such a mechanistic account. Within this account, learning is governed by three key quantities—surprise, belief uncertainty, and volatility [20, 21]. Surprise reflects the mismatch between expected and observed outcomes, signalling when beliefs need to be updated. Belief uncertainty represents the precision of current expectations, determining how strongly new information should influence updating. Volatility captures the tendency of which the environment itself is changing, indicating whether recent observations should be interpreted as noise or as evidence for a shift in underlying contingencies [4, 20, 22]. Together, these quantities capture distinct aspects of hierarchical belief updating in changing environments. Empirical work shows that subjective stress and physiological responses track these quantities during aversive learning, indicating that stress is shaped by inferred uncertainty rather than just objective uncertainty determined by task structure [6, 23, 24].

To test this framework, we applied the Volatile Kalman Filter (VKF), an analytical implementation of a hierarchical Bayesian model of stress that has been shown to effectively track learning under volatile environments [20]. The VKF was used to estimate trial-by-trial surprise, belief uncertainty, and volatility. Participants completed a probabilistic aversive learning task in which predictability and controllability were manipulated orthogonally. Subjective stress ratings, skin conductance responses, and reaction times were measured to characterise how model-derived uncertainty estimates related to subjective, physiological, and behavioural responses. We tested whether controllability modulates which computational uncertainty estimate—surprise, belief uncertainty, or volatility—best accounts for stress-related responses, and whether this relationship depends on predictability.

## Methods and Materials

### Participants

Thirty healthy adults (n = 30; 16 women, 14 men; age range = 18–40 years) were recruited through the University of Melbourne via online advertisements and campus flyers. Individuals with a history of major neurological or psychiatric disorders or a family history of epilepsy or other conditions that could interfere with the experimental procedures were excluded. Each participant completed two experimental conditions on separate days, one under controllable and one under uncontrollable conditions. All participants provided written informed consent prior to participation, and all procedures received approval from the university’s Human Research Ethics Committee. Participants were compensated at a rate of AUD $10 per 30 minutes.

### Probabilistic Aversive Learning Task

Participants first received instructions explaining the task (Figure. 1) and completed a brief practice session. Each condition consisted of 320 trials. On each trial (Figure. 1a), one of two rock-image cues was presented and participants predicted whether a snake would appear. The outcome then indicated whether a snake was present or absent, and each snake outcome coincided with an electric shock. In the controllable condition, correct predictions of a snake outcome were followed by a low-intensity shock, whereas incorrect predictions resulted in a high-intensity shock. In the uncontrollable condition, shock intensity was independent of prediction accuracy and was determined by the participant-specific shock sequence generated during the controllable session. Cue–outcome contingencies varied across ten unsignalled blocks (Figure. 1c). Snake outcomes occurred on 90% of trials in highly predictable blocks, 70% in moderately predictable blocks, and 50% in unpredictable blocks. The order of blocks was counterbalanced across participants. Full trial timing, shock calibration, and stimulation procedures are provided in the Supplement.

**Figure. 1.**
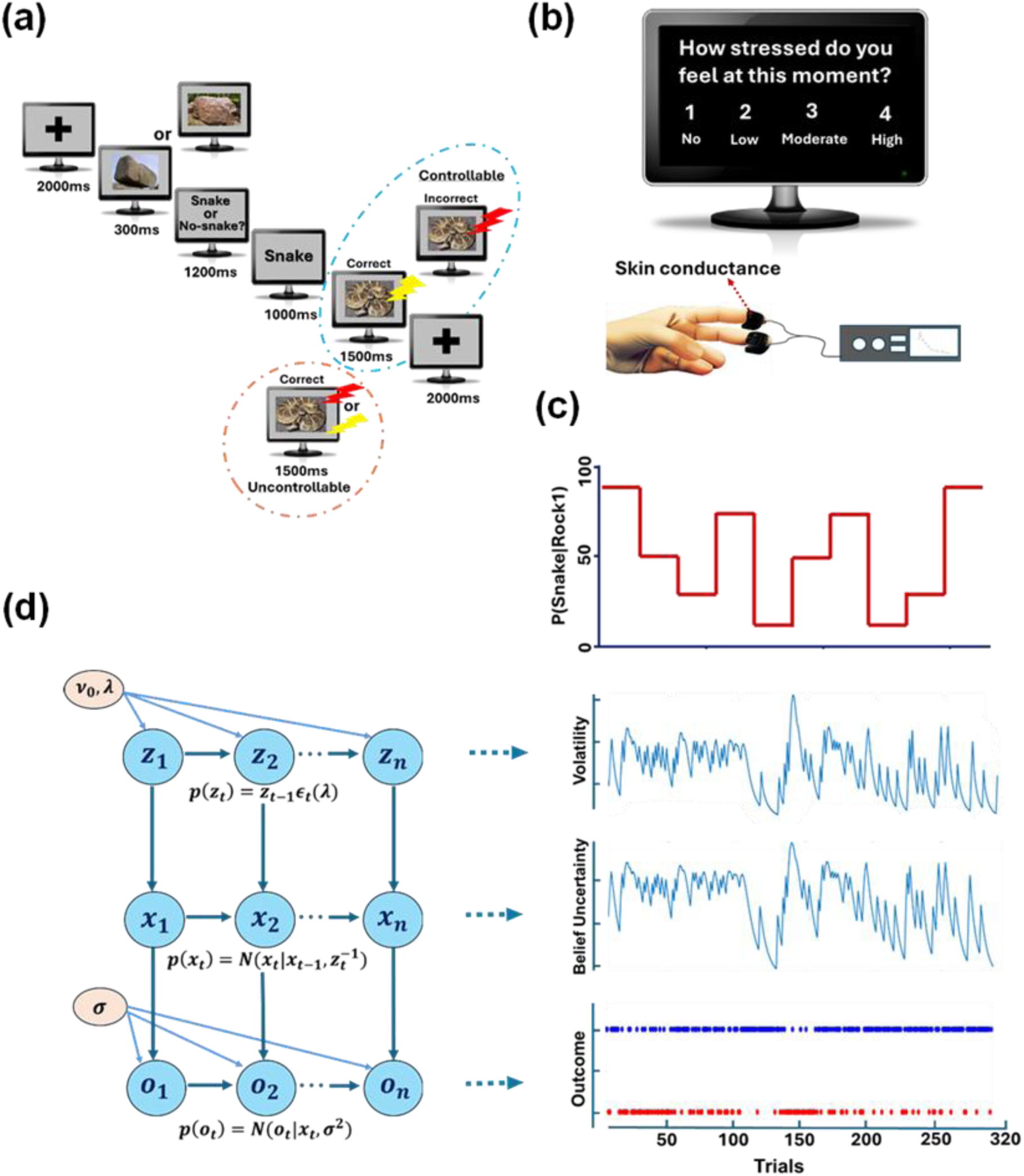
Task structure, experimental manipulations, stress measures, and computational model. **a**, Trial sequence and controllability manipulation. Each trial began with a fixation cross, followed by a rock cue that probabilistically predicted whether the outcome would be a snake or no snake. Participants made a binary prediction before the outcome was revealed. Snake outcomes were paired with electric shock. In the controllable condition, shock intensity depended on prediction accuracy on snake trials, with accurate predictions followed by low-intensity shocks and inaccurate predictions followed by high-intensity shocks. In the uncontrollable condition, shock intensity was independent of participants’ behaviour. **b**, Subjective and physiological measures. Subjective stress ratings were obtained every 4–6 trials using a rating scale, and skin conductance was recorded continuously from the fingers as an index of autonomic arousal. **c**, Predictability manipulation. Cue–outcome contingencies varied across unsignalled blocks, with highly predictable, moderately predictable, and unpredictable conditions. **d**, Volatile Kalman filter model. The model estimates the expected outcome *x_t_*, which represents the predicted probability of a snake, and environmental volatility *z_t_*, which reflects changes in cue–outcome relationships over time. The observed outcome *o_t_* represents the trial outcome ssnake or no snake). The parameter *σ*represents observation noise and captures variability in the outcomes. After each trial, beliefs about *x_t_* are updated based on the difference between the expected and observed outcome sprediction error). The magnitude of this update is determined by a trial-wise learning rate, which depends on belief uncertainty and inferred volatility. The volatility estimate *z_t_* is updated across trials according to the volatility update rate *λ*, with *v*_0_ specifying initial volatility.

### Subjective and Physiological Measures

Participants provided subjective stress ratings every 4–6 trials (Figure. 1b), totaling 65 ratings across 320 trials, using a five-point Likert scale ranging from 1 (not stressed) to 5 (high stress). Electrodermal activity was recorded continuously from the non-dominant hand as a measure of autonomic arousal. Full acquisition and preprocessing procedures are provided in the Supplement.

### Modelling of learning

Trial-by-trial learning was modelled using the Volatile Kalman Filter (VKF), a hierarchical Bayesian learning model designed to capture belief updating in environments with changing outcome contingencies [20]. The VKF extends the classical Kalman filter by jointly estimating outcome probabilities and environmental volatility, allowing the learning rate to adapt dynamically to inferred environmental stability. Model fitting was performed separately for each participant using Bayesian model inversion with a Laplace approximation to the posterior (Figure. 1d). Trial-wise surprise, belief uncertainty, volatility, and learning rate were subsequently derived from the fitted model. Full model equations and parameterisation are provided in the Supplement.

To investigate how latent learning processes influence stress responses, trial-wise surprise, belief uncertainty, and volatility were fitted in separate general linear models for each outcome measure. Correct prediction was included as a control predictor in all models to account for task performance effects, whereas models predicting stress ratings additionally included the previous trial’s rating to account for temporal dependence. Models were estimated separately for each participant using ordinary least-squares regression implemented in MATLAB (fitlm). Model fit was evaluated using the Bayesian Information Criterion, and values were entered into a random-effects Bayesian model comparison procedure implemented in the VBA toolbox [25, 26]. Full model specifications are provided in the Supplement.

### Statistical Analysis

All statistical analyses were performed in MATLAB (R2021b, MathWorks) using built-in functions and custom scripts developed for this study. Accuracy, reaction time, and skin conductance response amplitudes were averaged within each experimental condition and analysed separately using 2 × 3 repeated-measures ANOVAs with Controllability (controllable, uncontrollable) and Predictability (highly predictable, moderately predictable, unpredictable) as within-participant factors. Significant main effects and interactions were followed by Bonferroni-corrected pairwise comparisons. Regression coefficients obtained from participant-level general linear models were entered into group-level one-sample t-tests. Additional statistical and exclusion details are provided in the Supplement.

## Results

### Behavioural performance dissociates across predictability and controllability

A repeated-measures ANOVA on accuracy, defined as the proportion of trials on which participants correctly predicted the outcome, revealed significant main effects of controllability (F(1,29) = 17.19, p < 0.001) and predictability (F(2,58) = 86.31, p < 0.001), as well as a significant controllability × predictability interaction (F(2,58) = 3.52, p = 0.036). Accuracy was highest in highly predictable (HP) blocks and declined markedly in moderately predictable (MP) and unpredictable (UP) blocks under both controllable and uncontrollable conditions. Accuracy was significantly higher in the uncontrollable than controllable condition in HP blocks (p < 0.001) and MP blocks (p = 0.013), but not in UP blocks (p > 0.05). Thus, predictability exerted the strongest influence on performance overall, while the effect of controllability was evident only when in a highly predictable environment (Figure 2a).

**Figure. 2.**
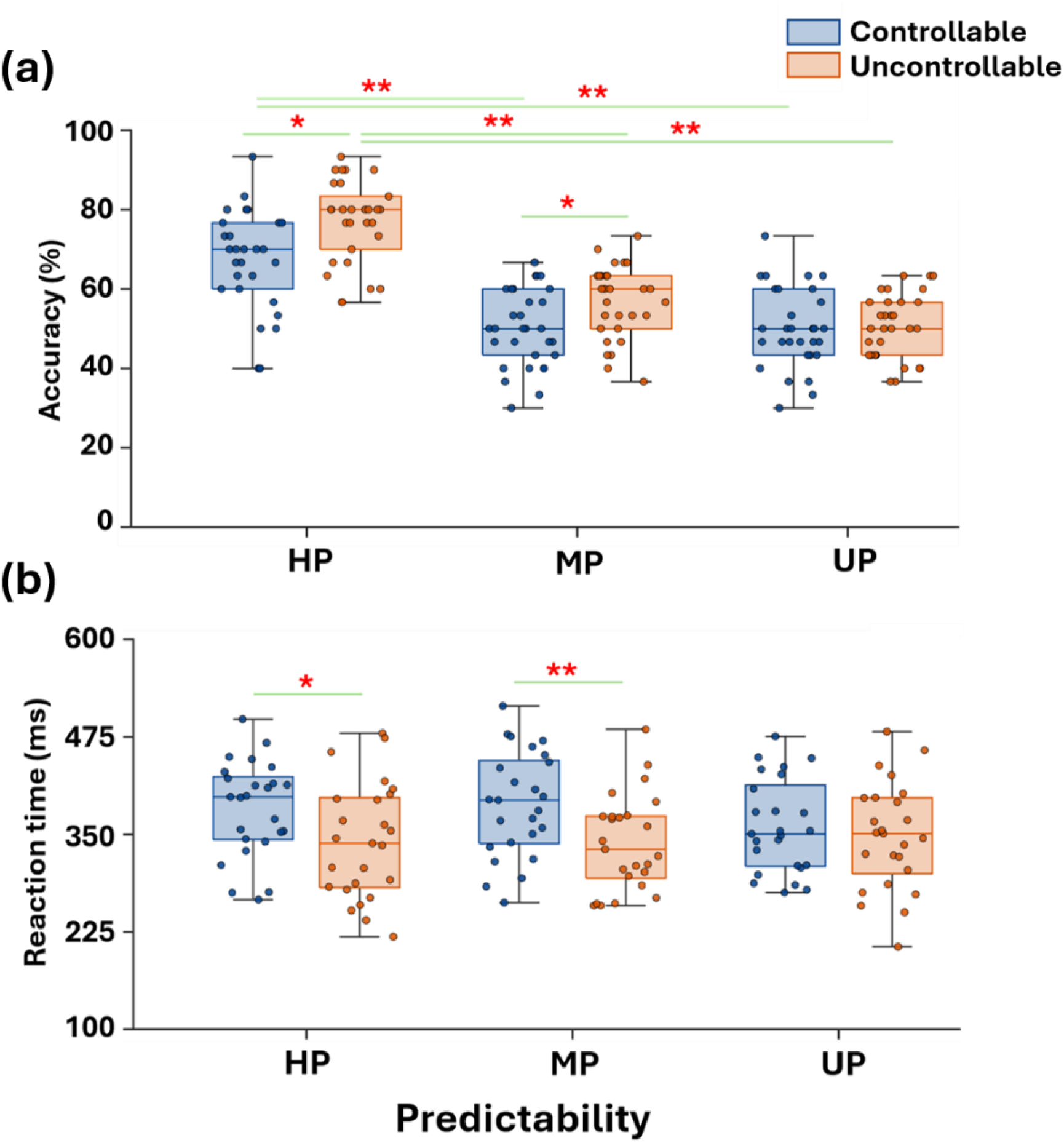
Behavioural accuracy and reaction time across controllability and predictability. (**a)** Prediction accuracy across highly predictable, moderately predictable and unpredictable blocks under controllable (blue) and uncontrollable (orange) conditions. Accuracy decreased with reduced predictability (main effect of predictability, *F*(2,58) = 86.31, *p* < 0.001) and was higher in the uncontrollable than controllable condition in HP and MP blocks (both *p* < 0.05), with no difference in UP blocks. (**b)** Reaction time across the same conditions and predictability levels. Responses were slower in the controllable than uncontrollable condition (main effect of controllability, *F*(1,24) = 13.34, *p* = 0.001), particularly in HP and MP blocks (both *p* < 0.05), with no difference in UP blocks. Points represent individual participants; boxplots show the median and interquartile range, with whiskers indicating the range.

Reaction time showed a distinct but related pattern (Figure 2b). A repeated-measures ANOVA revealed a significant main effect of controllability (F(1,24) = 13.34, p = 0.001) and a significant controllability × predictability interaction (F(2,48) = 3.34, p = 0.044), whereas the main effect of predictability was not significant (F(2,48) = 1.00, p > 0.05). Participants responded more slowly in the controllable than uncontrollable condition in HP blocks (p = 0.014) and MP blocks (p < 0.001), with no difference in UP blocks (p = 0.314). These findings indicate that predictability primarily determined how accurately participants learned the outcome structure, whereas controllability primarily influenced response speed when the environment was informative.

### Skin conductance responses reflect an interaction between predictability and controllability

In the controllable condition, peak skin conductance response (SCR) amplitudes were lower during HP and MP trials than during UP trials (Figure 3). In contrast, under uncontrollable conditions, SCRs remained lowest during HP trials, whereas MP and UP trials elicited indistinguishable responses. A 2 × 3 repeated-measures ANOVA confirmed significant main effects of controllability (F(1,174) = 25.12, p < 0.001) and predictability (F(2,174) = 48.49, p < 0.001), as well as a strong interaction (F(2,174) = 21.79, p < 0.001). Post hoc comparisons showed that, in the controllable condition, HP and MP did not differ (p > 0.05), whereas both were lower than UP sps ≤ 0.01). In the uncontrollable condition, HP responses were lower than both MP and UP (ps < 0.001), while MP and UP did not differ (p> 0.05).

**Figure. 3.**
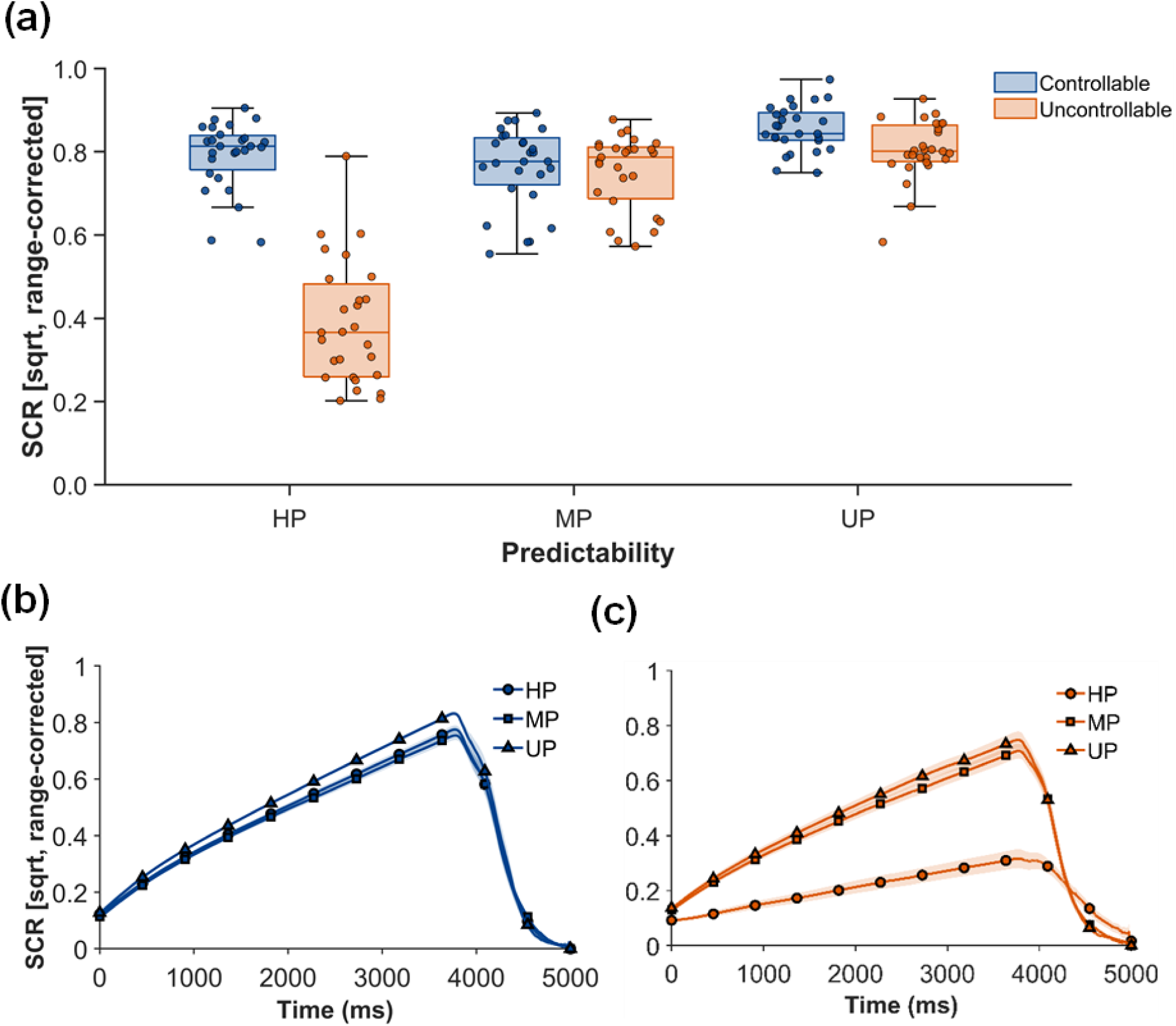
Skin conductance responses reflect an interaction between controllability and predictability. (a) Range-corrected, square-root-transformed skin conductance responses (SCRs) across highly predictable (HP), moderately predictable (MP), and unpredictable (UP) blocks under controllable (blue) and uncontrollable (orange) conditions. When controllable, SCRs were lower in HP and MP than in UP. When uncontrollable, SCRs in HP were reduced, whereas SCRs in MP and UP were similarly elevated. Points represent individual participants; boxplots show the median and interquartile range, with whiskers indicating the full range. (b) Time-resolved SCRs in the controllable condition for HP, MP, and UP blocks, showing lower SCRs in HP and MP than in UP across the trial. (c) Time-resolved SCRs in the uncontrollable condition for HP, MP, and UP blocks, showing reduced SCRs in HP and similarly elevated SCRs in MP and UP.

### Controllability modulates which uncertainty representations govern behavioural and autonomic responses

Model comparison conducted separately for subjective ratings, behavioural responses, and autonomic arousal revealed that controllability modulated which form of computational uncertainty best accounted for stress-related measures (Figure 4a). When outcomes were controllable, the volatility model showed the strongest evidence for subjective stress ratings (exceedance probability = 0.99), whereas belief uncertainty was strongly favoured during the uncontrollable condition (exceedance probability > 0.90). Reaction times models showed that belief uncertainty provided the best account of RT variability (exceedance probability = 0.92), whereas model evidence was distributed between surprise and volatility in the uncontrollable condition. For SCR variability, belief uncertainty provided the best account in the controllable condition (exceedance probability = 0.70), whereas volatility was favoured in the uncontrollable condition (exceedance probability = 0.72).

**Fig. 4.**
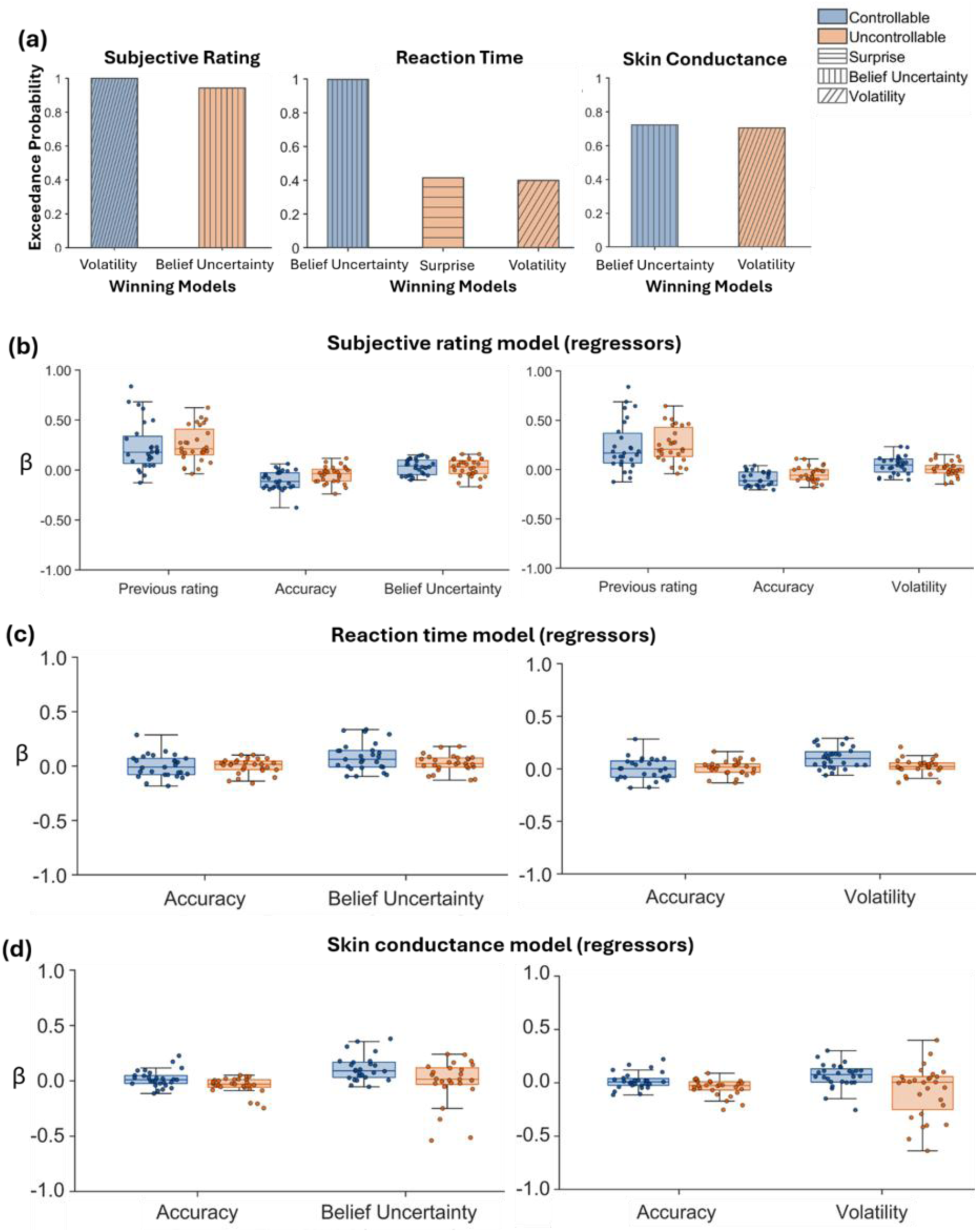
Controllability modulates which uncertainty representation explains subjective, behavioural and autonomic responses. (a) Exceedance probabilities for belief uncertainty, volatility and surprise models across subjective ratings, reaction time and skin conductance responses under controllable and uncontrollable conditions. Volatility was favoured for subjective ratings under control (exceedance probability = 0.99), whereas belief uncertainty was favoured under uncontrollable conditions (exceedance probability > 0.90). (b) Regression coefficients for subjective ratings showing effects of previous rating, belief uncertainty and volatility across conditions. Volatility predicted ratings under control, whereas belief uncertainty showed the strongest association under uncontrollable condition (ps < 0.05). (c) Regression coefficients for reaction time. Reaction times increased with uncertainty under control (ps ≤ 0.01), with reduced effects under uncontrollable condition. (d) Regression coefficients for skin conductance responses. Both belief uncertainty (p = 0.044) and volatility (p = 0.017) predicted responses under control, whereas effects were not reliable under uncontrollable conditions.

Regression analyses examined whether the computational uncertainty estimates explained trial-wise variations in subjective ratings, behavioural responses, and autonomic arousal. We additionally performed cross-modal analyses by correlating participant-level regression coefficients for belief uncertainty and volatility across response measures to determine whether these computational associations were shared. For subjective stress ratings (Fig. 4b), the previous rating strongly predicted the current report in both controllable (β = 0.23, p < 0.001) and uncontrollable (β = 0.25, p < 0.001) conditions, indicating substantial temporal dependence in subjective stress ratings. Volatility and belief uncertainty also contributed to trial-by-trial variations in subjective stress ratings, although both effects were significantly smaller than that of the previous rating. Under controllable conditions, volatility (β = 0.046, p < 0.01) and belief uncertainty (β = 0.033, p < 0.001) showed smaller effects than the previous rating. A similar pattern was observed under uncontrollable conditions, where both volatility (β = 0.009) and belief uncertainty (β = 0.022) also showed smaller effects than the previous rating sboth p < 0.001). Neither belief uncertainty nor volatility differed significantly between controllable and uncontrollable conditions. Reaction times showed associations with computational uncertainty estimates that depended on controllability (Fig. 4c). When outcomes were controllable, reaction times increased with both belief uncertainty (β = 0.085, p < 0.01) and volatility (β = 0.103, p < 0.001), indicating slower responses as environmental uncertainty increased. Under uncontrollable conditions, neither belief uncertainty (β = 0.025, p > 0.05) nor volatility (β = 0.021, p > 0.05) was associated with reaction times. SCR amplitudes showed weaker associations with these uncertainty estimates (Fig. 4d). In the controllable condition, SCR amplitudes increased with both belief uncertainty (β = 0.115, p < 0.001) and volatility (β = 0.068, p < 0.01). In contrast, neither belief uncertainty (β = −0.012, p > 0.05) nor volatility (β = −0.077, p > 0.05) was associated with SCR amplitudes under uncontrollable conditions. A paired t-test comparing participant-level β coefficients between conditions showed that the volatility β coefficients differed significantly, ts26) = 3.07, p < 0.01, whereas the corresponding difference in belief uncertainty β coefficients was not significant, t(27) = 1.79, p > 0.05. Cross-modal analyses showed that participant-level regression coefficients for belief uncertainty and volatility did not reliably covary across subjective ratings, reaction times, and skin conductance responses, indicating that uncertainty was represented differently across subjective, behavioural, and autonomic responses (see Supplementary Fig. S1).

### Learning rate and volatility updating show distinct associations with behavioural performance

Learning rate showed a robust negative association with prediction accuracy (Figure 5). Lower learning rates were associated with higher accuracy across predictability levels, with the strongest relationships in HP blocks in both the controllable sr = −0.92, p < 0.001) and uncontrollable conditions sr = −0.83, p < 0.001). The same relationship persisted in MP and UP blocks, although it became weaker as predictability declined. Learning rate showed no reliable association with reaction time. In contrast, volatility update rate showed condition-specific associations with behavioural performance (Figure 6): Higher volatility update rates were associated with greater accuracy only in the uncontrollable HP block (r = 0.39, p = 0.040) and controllable MP block (r = 0.49, p = 0.008). Higher volatility update rates were also associated with faster reaction times in the controllable MP condition sr = −0.41, p = 0.030), with no reliable associations in the uncontrollable condition.

**Figure. 5.**
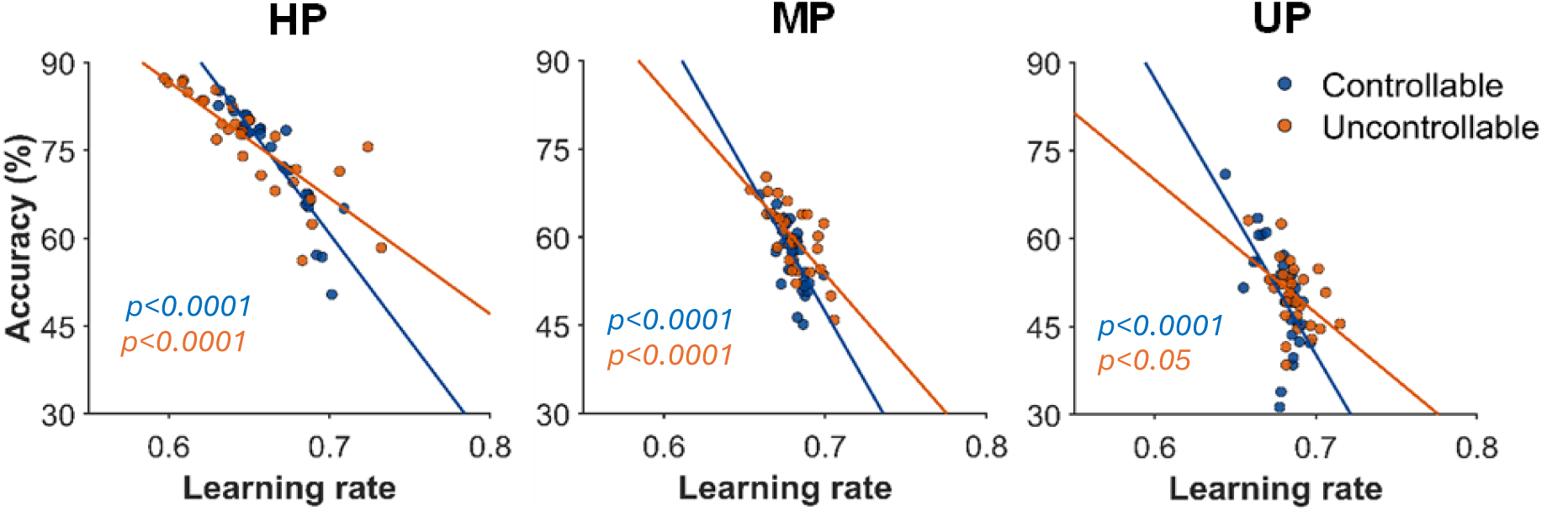
Learning rate is associated with accuracy. Relationship between learning rate and prediction accuracy across highly predictable, moderately predictable and unpredictable blocks under controllable (blue) and uncontrollable (orange) conditions. Accuracy decreased with increasing learning rate, with the strongest negative association in highly predictable blocks scontrollable r = −0.92; uncontrollable r = −0.83; both, p < 0.001).

**Figure 6.**
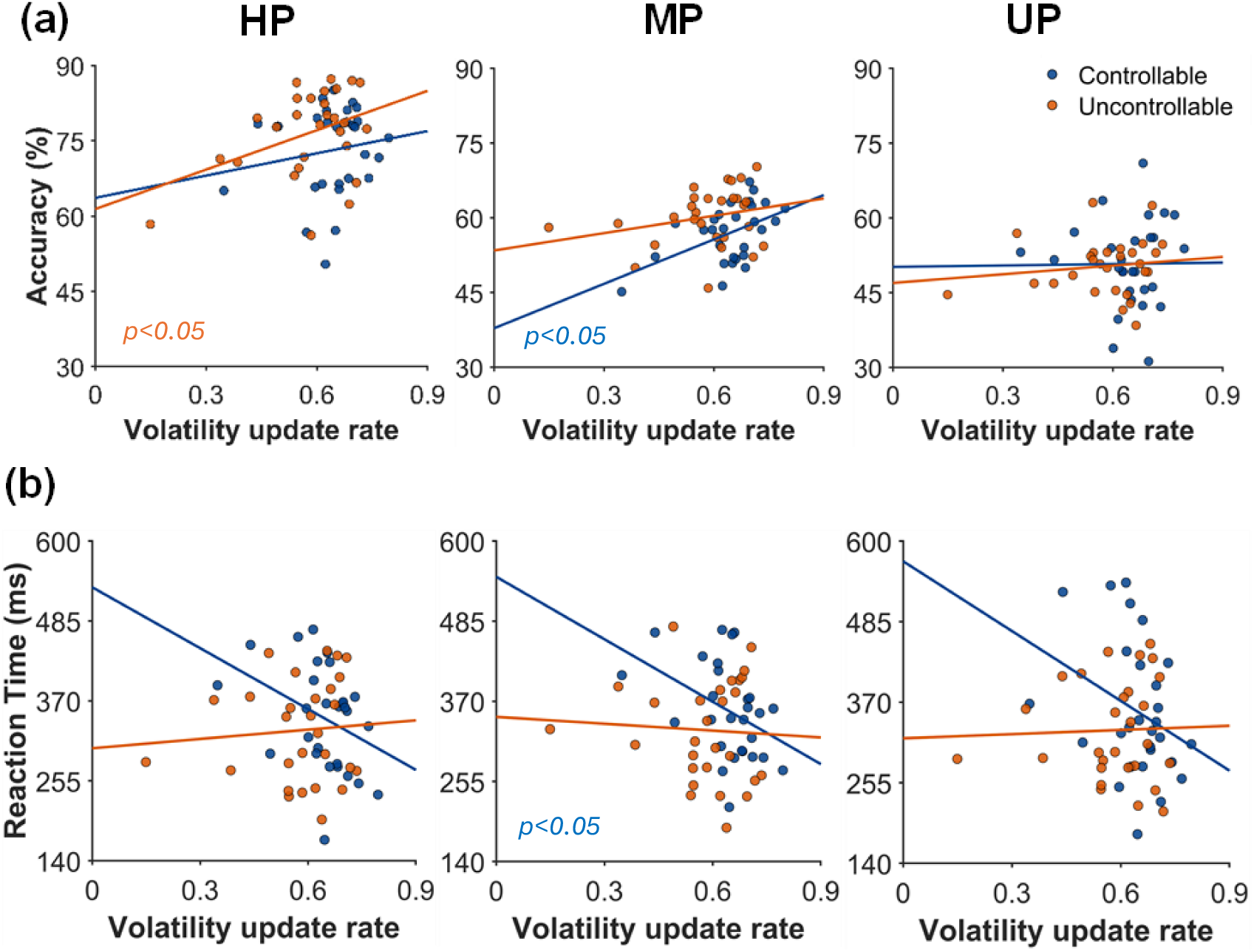
Volatility updating shows context-dependent associations with behaviour. **(a)** Relationship between volatility update rate and prediction accuracy across HP, MP and UP blocks under controllable sblue) and uncontrollable sorange) conditions. Positive associations were observed selectively, including in uncontrollable highly predictable blocks sr = 0.39, p = 0.040) and controllable moderately predictable blocks sr = 0.49, p = 0.008), with no consistent relationships across other conditions. **(b)** Relationship between volatility update rate and reaction time across the same predictability levels and conditions. Higher volatility update rates were associated with shorter reaction times in the controllable condition, reaching significance in moderately predictable blocks sr = −0.41, p = 0.030), with no reliable associations in the uncontrollable condition.

### Anxiety selectively predicts learning rate in predictable and controllable environments

Higher Beck Anxiety Inventory scores were associated with higher learning rates in HP (r = 0.47, p = 0.0088) and MP blocks (r = 0.48, p = 0.0091) under controllable condition (Figure 7). This relationship was not present in unpredictable blocks and was not observed at any predictability level in the uncontrollable condition. Volatility update rate was not significantly related to anxiety in either condition, and no significant associations were observed between depression or intolerance of uncertainty and any computational parameter. Detailed parameter associations are reported in the Supplementary.

**Figure. 7.**
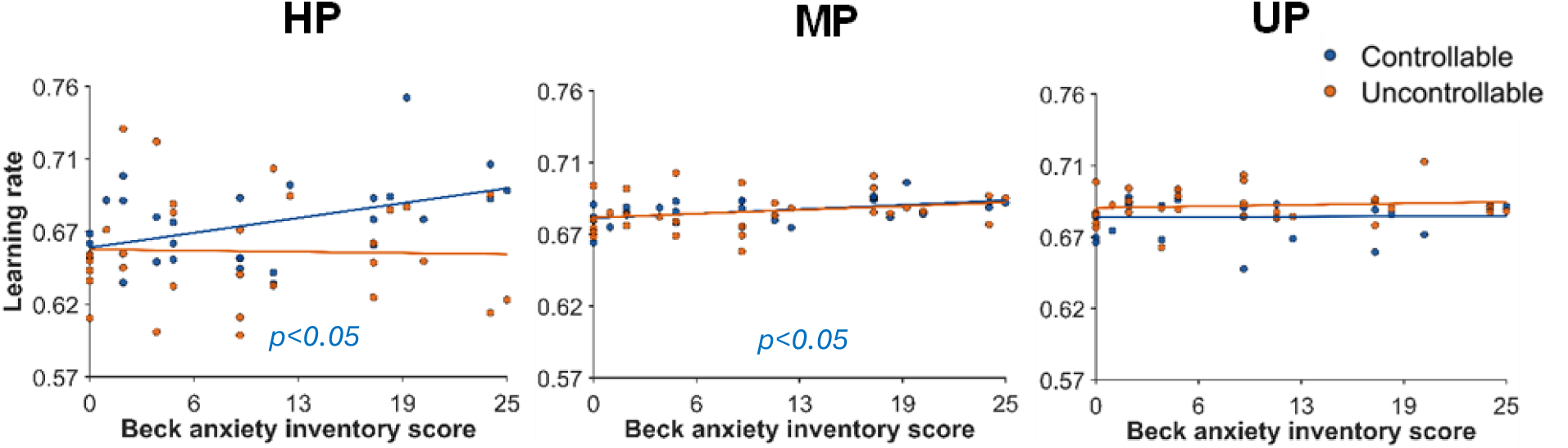
Anxiety selectively relates to learning rate under controllable conditions. Relationship between Beck Anxiety Inventory scores and learning rate across highly predictable (HP), moderately predictable (MP) and unpredictable (UP) blocks under controllable (blue) and uncontrollable (orange) conditions. Higher anxiety was associated with higher learning rates in the controllable highly predictable (r = 0.47, p = 0.0088) and moderately predictable blocks (r = 0.48, p = 0.0091), with no association in unpredictable blocks or in any uncontrollable condition.

## Discussion

The present study examined how predictability and controllability jointly shape behavioural, subjective, and autonomic responses during probabilistic aversive learning. The findings indicate that predictability was associated with prediction accuracy, whereas controllability was associated with reaction time and skin conductance responses. Model-based analyses further showed that controllability modulated which uncertainty estimates best accounted for subjective ratings, reaction times, and skin conductance responses. Together, these findings indicate that stress responses during aversive learning reflect both the predictability of environmental contingencies and the extent to which individuals can control shock intensity through accurate predictions.

Behavioural performance was strongly constrained by environmental predictability. Accuracy peaked under stable cue–outcome contingencies and declined as predictability decreased; reflecting that stable contingencies support precise expectations while less predictable ones provide unreliable information [20, 27, 28]. Controllability also influenced performance. Accuracy was higher in the uncontrollable than controllable condition in highly and moderately predictable blocks, but not in unpredictable blocks. This suggests that the uncontrollable condition placed fewer cognitive demands on outcome prediction. When shock intensity depended on accuracy, participants had to estimate cue–outcome contingencies while also considering whether their predictions could reduce shock intensity. This additional action–outcome component may have interfered with cue–outcome learning and reduced accuracy. This interpretation aligns with accounts proposing that good performances in controllable environments require integration of cue–outcome and action–outcome information, whereas uncontrollable environments allow predictions to rely more directly on environmental contingencies [12]. The absence of controllability effects in unpredictable blocks likely reflects the weak predictive value of cues, limiting performance in both conditions [20, 28].

For Skin conductance responses, controllability determined whether physiological stress reduced in MP blocks: MP block responses resembled UP blocks when accurate predictions allowed shock reduction, but resembled HP blocks when performance could not influence shock (Figure 3), consistent with prior work showing that controllable stress engages regulatory mechanisms that attenuate autonomic arousal [8, 29]. Computational modelling further clarified this pattern. SCRs were best explained by belief uncertainty when outcomes were controllable, indicating greater certainty reduced autonomic arousal when predictions could reduce shock intensity. This is consistent with evidence that physiological responses during probabilistic threat learning are related to uncertainty about expected outcomes [6] and that SCRs are larger when aversive outcomes are less expected [30]. When outcomes could not be influenced by behaviour, SCRs were best explained by volatility, indicating that autonomic arousal was more closely related to inferred changes in cue–outcome contingencies. This is consistent with evidence that lack of control alters behavioural adjustment [8] and is associated with stronger expectations that environmental contingencies may change [14].

Reaction-time dynamics provided additional evidence that controllability modulates how computational uncertainty shapes behavioural responses. Participants responded more slowly in the controllable condition during HP and MP blocks, suggesting greater cognitive engagement during response selection when cue–outcome contingencies can be used to mitigate aversive outcomes [12, 31]. Model comparison showed that belief uncertainty explained reaction-time variability when shock intensity depended on prediction accuracy, whereas no single computational estimate explained reaction time when shock intensity was independent of participants’ predictions. Under controllable conditions, uncertainty about expected outcomes mattered to predictions participants made to reduce shock intensity. Greater belief uncertainty therefore likely increased information gathered and reaction time before responding, consistent with evidence of increased uncertainty prolonged reaction time [6, 32, 33].

Cross-modal analyses examined whether individual differences in the effects of belief uncertainty and volatility were shared across response domains. Participant-level effects were not reliably correlated across subjective ratings, reaction times, and skin conductance responses, indicating that participants who showed a stronger computational effect in one response domain did not necessarily show a similarly strong effect in another. This limited cross-domain correspondence is consistent with evidence that subjective experience, behaviour, and physiological arousal are often only partially aligned [34–36], and with the proposal that conscious threat-related experiences and behavioural or physiological threat responses arise from partly distinct systems [37].

We tested whether learning rate and volatility updating in the VKF models correlated with behaviours. Lower learning rates were associated with higher accuracy, with the strongest effect in highly predictable environments. The result is consistent with Bayesian learning accounts, which predicts that learning rates should be lower when environmental contingencies are stable because unexpected outcomes are more likely to reflect stochastic noise than genuine changes in states [4, 5, 23]. This relationship was similar in both controllable and uncontrollable conditions, indicating that learning rate primarily reflects adaptation to outcome contingencies rather than control over outcomes. Learning rate showed no association with reaction time, suggesting that the extent of belief updating was related to prediction accuracy but did not determine how quickly participants translated those beliefs into a response. This is consistent with decision-making accounts in which reaction time reflects evidence accumulation and response selection [38, 39].

Volatility updating had a more selective effect on behaviour than learning rate. Higher volatility update rates were associated with greater accuracy only in HP blocks in the uncontrollable condition and in MP blocks in the controllable condition. This pattern suggests that volatility updating was most useful when cue–outcome probabilities provided reliable information about changes in contingencies, consistent with previous accounts of volatility-guided learning [40, 41]. Higher volatility update rates were also selectively associated with faster responses in the controllable condition, particularly in MP blocks, where tracking changes in cue–outcome contingencies were behaviourally relevant because accurate predictions could reduce shock intensity [8, 42].

Higher Beck Anxiety Inventory scores were associated with increased learning rates in highly and moderately predictable blocks when outcomes were controllable. This pattern indicates that anxiety was linked to heightened trial-by-trial belief updating even when environmental structure was stable and action–outcome contingencies were present. Under normative Bayesian accounts, stable environments favour lower learning rates to prevent overfitting to stochastic fluctuations [4, 5, 40]. Elevated learning rates in more anxious individuals may therefore reflect oversensitivity to recent outcomes despite stable structure. This interpretation aligned with computational evidence that anxiety reduces flexibility in adjusting learning rates based on environmental stability and impairs discrimination between stable and changing environments, resulting in less effective adaptation [43–45]. No reliable relationships were observed between anxiety and volatility updating, depression, or intolerance of uncertainty.

A key limitation is that the controllable and uncontrollable sessions were not counterbalanced. The yoked design required this fixed order, so the uncontrollable session replayed the participant-specific controllable session shock sequence, matching shock timing and intensity. Matching shock exposure was important to minimise the possibility that differences between conditions were driven by differences in the aversive stimulation received, rather than by controllability. Counterbalancing would have required a predefined sequence or a sequence generated from another participant, thereby sacrificed this participant-specific matching. Nevertheless, because the uncontrollable session always followed the controllable session, the higher accuracy and faster reaction times observed in this condition may partly reflect practice or familiarity effects. However, these differences were not consistently observed across predictability levels, and skin conductance responses showed condition-dependent patterns rather than a uniform reduction in the second session, suggesting that session order alone is unlikely to explain the findings.

In conclusion, the present findings indicate that stress responses during aversive learning reflect hierarchical inference about environmental contingencies rather than uniform reactions to aversive threat. Predictability influenced the reliability of cue–outcome learning and the stability of belief updating, whereas controllability determined which computational representations of uncertainty were most strongly associated with subjective, behavioural, and physiological responses. This double dissociation suggests that whether behavioural predictions can influence aversive outcomes does not simply alter the magnitude of stress responses, but changes how uncertainty is represented. The absence of reliable cross-domain coupling further indicates that subjective ratings, behavioural responses, and autonomic arousal reflect different computational representations of uncertainty. Collectively, these findings support a computational account of stress in which responses are jointly shaped by the predictability of environmental contingencies and the extent to which behaviour can influence threat outcomes.

## Acknowledgments

We thank Katharina Wellstein for valuable discussions on hierarchical Bayesian modelling, which guided the selection of the modelling framework adopted in this study. We are grateful to Maya Schenker for training in the use of the ADInstruments PowerLab and LabChart data acquisition system and to Linzhi Tao for technical guidance on the Spartan high-performance computing platform. This project was funded by the 2023 Melbourne School of Psychological Sciences Research Incentive Scheme. Daniyal Rajput was supported by an Australian Government Research Training Program Scholarship.

## Data and Code Availability

The anonymised data supporting the findings are available on Figshare (Rajput et al., 2026). Analysis code for data processing, statistical analysis, and figure generation is available at https://github.com/daniyalrajput/predictability-controllability-aversive-learning.git.

Computational modelling code is available at https://github.com/daniyalrajput/stress_study_KF_VKF_scripts.git.

