## Supplementary File for "Predictability and controllability shape aversive learning and stress responses through independent computational mechanisms"

### Supplemental Methods

#### Participants and Procedure

Thirty healthy adults (16 women, 14 men; age range, 18–40 years) participated in the study. Full recruitment, exclusion, consent, and ethical approval details are provided in the main manuscript. Participants first received task instructions and completed a 30-trial practice session to confirm task understanding and familiarity with the subjective stress rating scale. The controllable session was completed first. The uncontrollable session used the participant-specific sequence of low- and high-intensity shocks generated during the controllable session, allowing shock timing and intensity exposure to be closely matched within participants.

#### Task Procedure and Experimental Manipulations

Each session consisted of 320 trials. A trial began with one of two rock-image cues presented for 300ms (±50ms). Participants predicted whether a snake would appear by pressing the left or right arrow key. The left arrow corresponded to snake and the right arrow to no snake, with the mapping held constant across participants. Responses were made within 1200ms. The selected prediction was displayed for 1000ms (±200ms), followed by the outcome for 1500ms (±200ms). Snake outcomes coincided with electric shock. Each trial ended with a 2000ms fixation interval.

Cue-outcome contingencies varied across 10 unsignalled blocks of 25-35 trials. The task contained four highly predictable blocks, four moderately predictable blocks, and two unpredictable blocks. Block order was counterbalanced, and participants were told that probabilities would change and might sometimes appear random.

### Shock Calibration and Stimulation

Electric shocks were delivered using a Stimulus Isolator controlled through customised LabChart software and a PowerLab interface (ADInstruments, Dunedin, New Zealand). Stimulation was administered through an electrode positioned adjacent to the first dorsal interosseous muscle of the left hand.

Before the controllable and uncontrollable sessions, shock intensity was calibrated separately using a staircase procedure to account for individual differences in pain sensitivity (Gracely et al., 1988). Shock intensity was gradually increased (0.1–8 mA) across up to 80 trials, and after each stimulation participants rated the perceived intensity on a four-point scale ranging from low to painful (1 = low, 2 = mild, 3 = uncomfortable, 4 = painful). Calibration continued until participants consistently identified the stimulation level corresponding to the boundary between uncomfortable and painful (3/4 boundary). This value was defined as the high-intensity shock level for that session. The low-intensity shock was set at 40% of this value, resulting in a weaker but clearly perceptible stimulus. This procedure ensured that stimulation remained individually tolerable while allowing participants to reliably distinguish between low- and high-intensity shocks across conditions.

### Subjective and Physiological Measures

Electrodermal activity was recorded continuously at 1000 Hz using two electrodes attached to the intermediate phalanges of the index and third fingers of the non-dominant hand. Skin conductance signals were acquired using an FE116 GSR Amp controlled through customised LabChart software via a PowerLab interface (ADInstruments, Dunedin, New Zealand).

**Skin Conductance Preprocessing**

Skin conductance data were preprocessed using the PsychoPhysiological Modelling toolbox [version 5.0; [1]], following standard procedures for skin conductance analysis [2]. Data were inspected for signal dropout, excessive noise, unstable baselines, poor electrode contact, and participant movement. Signal dropout was defined as prolonged near-zero conductance (≤0.05 μS).

Movement-related artefacts were identified using a custom MATLAB script that computed the second derivative of the signal to detect rapid changes. Samples in which the second derivative exceeded three standard deviations from the participant-level mean were classified as artefactual and replaced by linear interpolation between adjacent valid samples. Trials with substantial artefacts or missing data were excluded, and participants with a high proportion of these trials were excluded from subsequent analyses.

Following artefact correction, signals were band-pass filtered using a first-order Butterworth filter (0.05–5 Hz) to remove slow baseline drift and high-frequency noise while preserving phasic skin conductance responses. Filtered signals were square-root transformed to reduce skewness and range-corrected within participant to account for individual differences in skin conductance amplitude [3]. Anticipatory skin conductance responses were defined as the peak phasic response within the 2-second pre-outcome window on each trial. Peak amplitudes were normalised within participants to allow comparisons across individuals and experimental conditions.

**Volatile Kalman Filter**

Trial-wise learning was modelled using the Volatile Kalman Filter (VKF; Fig. 1e), a hierarchical Bayesian model for environments with changing outcome contingencies [4]. The VKF jointly estimates expected binary outcomes and environmental volatility, allowing learning rate to adapt to inferred environmental stability. The model was implemented using the publicly available MATLAB implementation provided by [4].

The VKF was applied to all trials. Outcomes were coded as a binary variable reflecting shock intensity with low-intensity shock coded as 0 and high-intensity shock coded as 1. This specification defines the observed outcome $o_t$ at each trial and allows the model to track trial-by-trial learning.

At the first level, the VKF tracks beliefs about the probability of outcomes. Beliefs are represented by a latent mean $m_t$, expressed in log-odds space and mapped to probabilities using the logistic sigmoid function $s(\cdot)$. On each trial, beliefs are updated according to a precision-weighted prediction error, defined as the difference between the observed outcome and the predicted probability of shock.

At the first level of the model, the observed outcome $o_t$ is represented. At the second level, beliefs about outcome probability are represented by a latent mean $m_t$, expressed in log-odds space and mapped to probabilities using the logistic sigmoid function $s(\cdot)$. Beliefs are updated on each trial according to

$$m_t = m_{t-1} + k_t(o_t - s(m_{t-1}))$$

where $k_t$ denotes the trial-wise learning rate and $o_t - s(m_{t-1})$ denotes the prediction error. Belief uncertainty $w_t$ determines the influence of new observations on belief updating and evolves according to

$$w_t = (1 - k_t)(w_{t-1} + v_{t-1})$$

where $w_{t-1}$ denotes belief uncertainty on the previous trial and $v_{t-1}$ denotes environmental volatility on the previous trial.

At the third level of the model, environmental volatility $v_t$ captures inferred changes in outcome contingencies over time and is updated according to

$$v_t = v_{t-1} + \lambda((m_t - m_{t-1})^2 + w_{t-1} + w_t - 2w_{t-1,t} - v_{t-1})$$

where $\lambda$ is the volatility update rate and $w_{t-1,t}$ denotes the lag-one covariance. Increases in estimated volatility increase subsequent learning rates, allowing the model to adapt more rapidly when contingencies change.

Model fitting was performed separately for each participant using Bayesian model inversion with a Laplace approximation to the posterior, as implemented in the CBM toolbox. Three participant-level parameters were estimated: the volatility update rate $\lambda$, the initial volatility $v_0$, and the observation noise parameter $\omega$. Trial-wise latent trajectories $(m_t, w_t, v_t, k_t)$ were subsequently derived from the fitted model. Surprise was defined as the prediction error, $o_t - s(m_{t-1})$. These model-derived quantities were then included as regressors in subsequent analyses of behavioural and physiological responses.

**Modelling of Stress Responses**

Separate general linear models were estimated for subjective stress ratings, reaction times, and skin conductance responses under each controllability condition. All predictors were z-scored within participant before model estimation to allow comparison of regression coefficients across regressors and individuals. The regression models were defined as

$$\text{Rating}^{(k)} = \beta_1 \cdot \text{Rating}^{(k-1)} + \sum_{i(k-1)}^{i(k)} \beta_2 \cdot \text{CorrectPrediction}^{(k)} + \beta_3 \cdot \text{ModelTrajectory}^{(k)} + \varepsilon^{(k)}$$

$$\text{SCR or RT}^{(k)} = \beta_1 \cdot \text{CorrectPrediction}^{(k)} + \beta_2 \cdot \text{ModelTrajectory}^{(k)} + \varepsilon^{(k)}$$

where $k$ denotes the trial number, $\beta$ represents regression coefficients, and $\epsilon_k$ denotes the residual error term. $ModelTrajectory_k$ corresponds to one of the VKF-derived signals (surprise, belief uncertainty, or volatility). All predictors were z-scored within participant prior to model estimation to allow comparison of regression coefficients across individuals and regressors.

Models were estimated separately for each participant using ordinary least-squares regression implemented in MATLAB (fitlm). Separate Bayesian model comparisons were conducted for subjective stress ratings, skin conductance responses, and reaction times. Within each response domain, three competing models were compared, with surprise, belief uncertainty, or volatility entered separately as the model-derived predictor. Thus, model comparison tested which computational estimate best explained trial-by-trial variation in each subjective, behavioural, and physiological response measure. For subjective stress ratings, previous trial rating and prediction accuracy were included as covariates, whereas prediction accuracy was included as a covariate for skin conductance responses and reaction times.

### Statistical Analysis

Accuracy, reaction time, and SCR amplitudes were averaged within each condition and analysed separately using 2 x 3 repeated-measures analyses of variance with Controllability and Predictability as within-participant factors. Significant effects were followed by Bonferroni-corrected comparisons. Values exceeding 3 standard deviations from the relevant condition mean were excluded.

Participant-level regression coefficients were entered into group-level one-sample tests. Pearson correlations assessed associations between VKF parameters and behavioural or questionnaire measures within each predictability and controllability condition. Cross-domain correlations compared participant-level belief-uncertainty and volatility coefficients across subjective ratings, reaction times, and SCRs.

### Supplemental Results

### Detailed Behavioural and SCR Results

Detailed post hoc comparisons are reported in Table S1. In the controllable condition, HP accuracy exceeded MP and UP accuracy (both $p < 0.001$), whereas MP and UP did not differ ($p > 0.05$). The same pattern was observed in the uncontrollable condition, with HP exceeding MP and UP (both $p < 0.001$), whereas MP and UP did not differ ($p > 0.05$).

Detailed post hoc comparisons are reported in Table S1. Reaction time did not differ significantly across predictability levels within either controllability condition. In the controllable condition, the comparison between moderately predictable and unpredictable blocks showed a trend toward slower responses in moderately predictable blocks ($p = .070$).

### Cross-Domain Coupling

Belief-uncertainty coefficients from subjective-rating models were not correlated with SCR coefficients under control ($r = 0.159$, $p >0.05$) or without control ($r = 0.041$, $p > 0.05$). Volatility coefficients were likewise unrelated across ratings and SCR under control ($r = 0.089$, $p > 0.05$) and without control ($r = 0.295$, $p >0.05$).

Rating-reaction-time correlations were also nonsignificant. Belief uncertainty showed no relationship under control ($r = 0.224$, $p >0.05$) or without control ($r = -0.127$, $p >0.05$). Volatility showed no reliable relationship under control ($r = 0.342$, $p >0.05$) or without control ($r = .144$, $p >0.05$). These results indicate limited participant-level coupling of uncertainty effects across response domains (Figure S1).

### Learning Rate and Volatility Updating

Detailed associations are shown in Figure S2 and Table S5. Significant associations with accuracy were observed in the uncontrollable HP and controllable MP conditions, as reported in the main text. For reaction time, the significant association in controllable MP blocks was accompanied by similar negative trends in controllable HP ($r = -0.37$, $p = 0.054$) and UP blocks ($r = -0.32$, $p = 0.092$). No reliable reaction-time associations were observed in the uncontrollable condition.

### Affective Traits and Model Parameters

Higher Beck Anxiety Inventory scores were associated with higher learning rates in controllable highly predictable and moderately predictable blocks. No association was present in controllable unpredictable blocks or any uncontrollable block. Volatility update rate was not significantly related to anxiety, although a negative trend occurred under control. Depression and intolerance of uncertainty were not significantly associated with learning rate or volatility update rate.

Detailed associations are reported in Table S5. Anxiety was not associated with learning rate in the controllable unpredictable condition ($r = 0.02$, $p > 0.05$) or in the uncontrollable highly predictable ($r = -0.03$, $p > 0.05$), moderately predictable ($r = 0.25$, $p > 0.05$), or unpredictable blocks ($r = 0.11$, $p > 0.05$). Volatility update rate was not significantly related to anxiety in either condition, although a negative trend was observed in the controllable condition ($r = -0.34$, $p > 0.05$) and was weaker in the uncontrollable condition ($r = -0.15$, $p > 0.05$). No significant associations were observed between depression or intolerance of uncertainty scores and any computational parameter across predictability or controllability conditions.

### Supplemental Tables

#### Table S1. Behavioural Analyses of Variance and Key Pairwise Comparisons

| Measure | Effect / Comparison | Statistic | p Value |
|---|---|---|---|
| Accuracy | Controllability | F(1,29) = 17.19 | < .001 |
| Accuracy | Predictability | F(2,58) = 86.31 | < .001 |
| Accuracy | Controllability x Predictability | F(2,58) = 3.52 | .036 |
| Accuracy | HP: uncontrollable vs controllable | - | < .001 |
| Accuracy | MP: uncontrollable vs controllable | - | .013 |
| Accuracy | UP: uncontrollable vs controllable | - | .784 |
| Reaction time | Controllability | F(1,24) = 13.34 | .001 |
| Reaction time | Predictability | F(2,48) = 1.00 | .376 |
| Reaction time | Controllability x Predictability | F(2,48) = 3.34 | .044 |
| Reaction time | HP: controllable vs uncontrollable | - | .014 |
| Reaction time | MP: controllable vs uncontrollable | - | < .001 |
| Reaction time | UP: controllable vs uncontrollable | - | .314 |

#### Table S2. Skin Conductance Analyses of Variance and Pairwise Comparisons

| Effect / Comparison | Statistic | p Value |
|---|---|---|
| Controllability | F(1,174) = 25.12 | < .001 |
| Predictability | F(2,174) = 48.49 | < .001 |
| Controllability x Predictability | F(2,174) = 21.79 | < .001 |
| Controllable: HP vs MP | - | .96 |
| Controllable: HP/MP vs UP | - | <= .01 |
| Uncontrollable: HP vs MP/UP | - | < .001 |
| Uncontrollable: MP vs UP | - | .81 |

#### Table S3. Random-Effects Bayesian Model Comparison

| Response | Condition | Favoured Model | Exceedance Probability |
|---|---|---|---|
| Subjective rating | Controllable | Volatility | .99 |
| Subjective rating | Uncontrollable | Belief uncertainty | > .90 |
| Reaction time | Controllable | Belief uncertainty | .92 |
| Reaction time | Uncontrollable | Surprise / volatility | .41 / .40 |
| Skin conductance | Controllable | Belief uncertainty | .70 |
| Skin conductance | Uncontrollable | Volatility | .72 |

### Table S4. Participant-Level Regression Coefficients

| Response | Predictor | Controllable beta (p) | Uncontrollable beta (p) |
|---|---|---|---|
| Rating | Previous rating | .230 (< .001) | .250 (< .001) |
| Rating | Belief uncertainty | .033 (< .001) | .022 (< .001) |
| Rating | Volatility | .046 (< .01) | .009 (< .001) |
| Reaction time | Belief uncertainty | .085 (< .01) | .025 (> .05) |
| Reaction time | Volatility | .103 (< .001) | .021 (> .05) |
| Skin conductance | Belief uncertainty | .115 (< .001) | -.012 (> .05) |
| Skin conductance | Volatility | .068 (< .01) | -.077 (> .05) |

### Table S5. Associations Between Computational Parameters, Behaviour, and Anxiety

| Association | Condition | Level | r | p |
|---|---|---|---|---|
| Learning rate / accuracy | Controllable | HP | -.92 | < .001 |
| Learning rate / accuracy | Uncontrollable | HP | -.83 | < .001 |
| Learning rate / accuracy | Controllable | MP | -.69 | < .001 |
| Learning rate / accuracy | Uncontrollable | MP | -.70 | < .001 |
| Learning rate / accuracy | Controllable | UP | -.65 | < .001 |
| Learning rate / accuracy | Uncontrollable | UP | -.46 | .011 |
| Volatility update / accuracy | Uncontrollable | HP | .39 | .040 |
| Volatility update / accuracy | Controllable | MP | .49 | .008 |
| Volatility update / reaction time | Controllable | HP | -.37 | .054 |
| Volatility update / reaction time | Controllable | MP | -.41 | .030 |
| Volatility update / reaction time | Controllable | UP | -.32 | .092 |
| Anxiety / learning rate | Controllable | HP | .47 | .0088 |
| Anxiety / learning rate | Controllable | MP | .48 | .0091 |
| Anxiety / learning rate | Controllable | UP | .02 | .906 |
| Anxiety / learning rate | Uncontrollable | HP | -.03 | .880 |
| Anxiety / learning rate | Uncontrollable | MP | .25 | .182 |
| Anxiety / learning rate | Uncontrollable | UP | .11 | .580 |

### Table S6. Cross-Domain Correlations of Computational Effects

| Trajectory | Response Pair | Condition | r | p |
|---|---|---|---|---|
| Belief uncertainty | Rating-SCR | Controllable | .159 | .419 |
| Belief uncertainty | Rating-SCR | Uncontrollable | .041 | .840 |
| Volatility | Rating-SCR | Controllable | .089 | .653 |
| Volatility | Rating-SCR | Uncontrollable | .295 | .128 |
| Belief uncertainty | Rating-RT | Controllable | .224 | .252 |
| Belief uncertainty | Rating-RT | Uncontrollable | -.127 | .520 |
| Volatility | Rating-RT | Controllable | .342 | .075 |
| Volatility | Rating-RT | Uncontrollable | .144 | .473 |

### Supplemental Figures

**Figure S1.** Limited cross-domain coupling of uncertainty effects. Participant-level belief-uncertainty and volatility coefficients from subjective-rating models were correlated with corresponding coefficients from skin conductance and reaction-time models under controllable and uncontrollable conditions. None of the correlations was statistically significant.

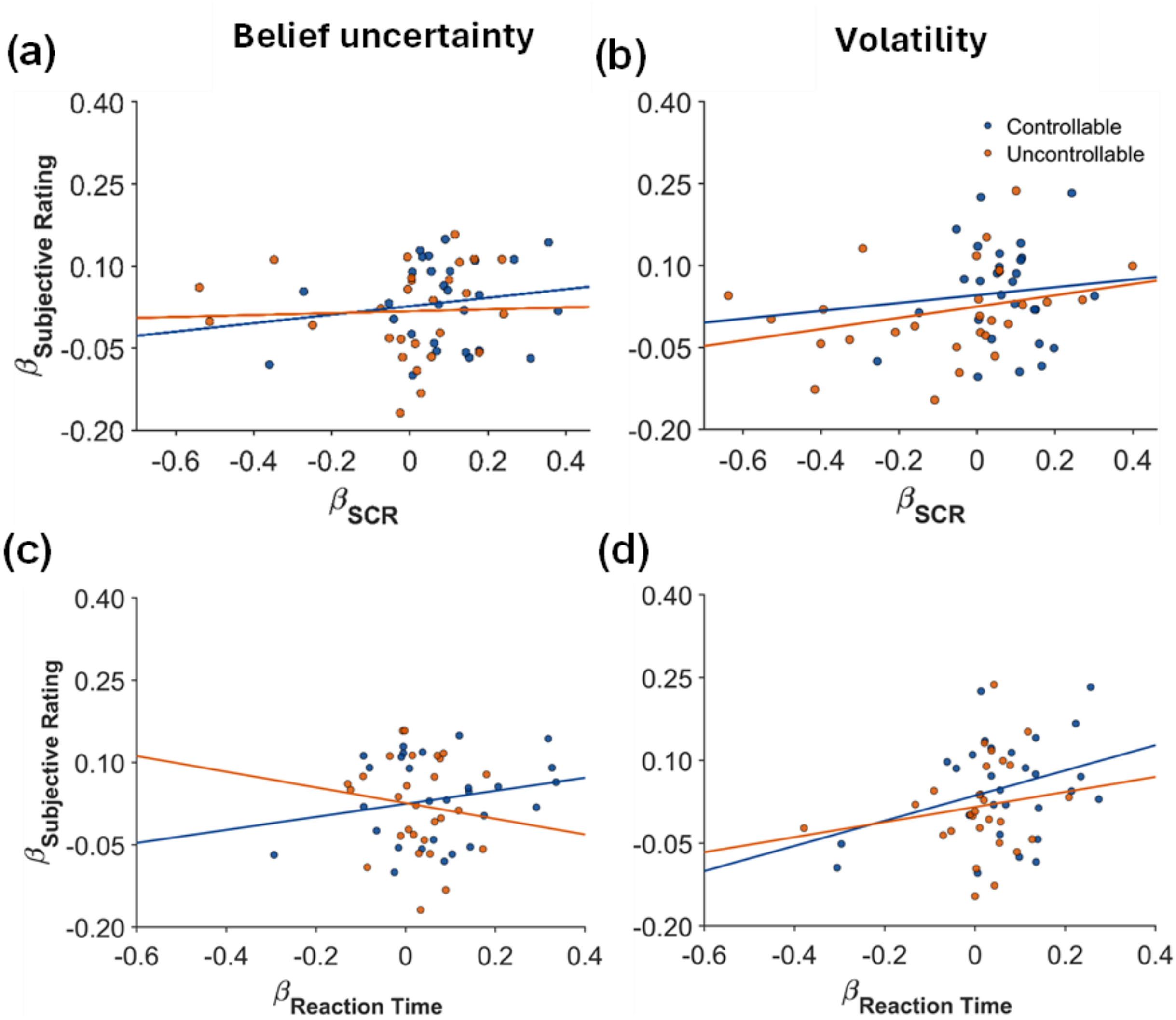

**Figure S2. Associations between volatility updating and behavioural performance across predictability and controllability. (a)** Relationships between volatility update rate and prediction accuracy across highly predictable (HP), moderately predictable (MP), and unpredictable (UP) blocks under controllable and uncontrollable conditions. Volatility update rate was positively associated with prediction accuracy in the uncontrollable condition during HP blocks ($r = 0.39$, $p = 0.040$) and in controllable condition during MP blocks ($r = 0.49$, $p = 0.008$). The remaining associations were not significant: controllable HP, $r = 0.16$, $p>0.05$; uncontrollable MP, $r = 0.26$, $p > 0.05$; controllable UP, $r = 0.01$, $p>0.05$; and uncontrollable UP, $r = 0.13$, $p>0.05$. **(b)** Relationships between volatility update rate and reaction time across the same predictability blocks and controllability conditions. In the controllable condition, higher volatility update rates were associated with shorter reaction times during MP blocks ($r = -0.41$, $p = 0.030$), with similar but nonsignificant associations during HP ($r = -0.37$, $p = 0.054$) and UP blocks ($r = -0.32$, $p = 0.092$). In uncontrollable conditions, associations were not significant during HP ($r = .08$, $p > 0.05$), MP ($r = -0.06$, $p > 0.05$), or UP blocks ($r = 0.04$, $p > 0.05$).

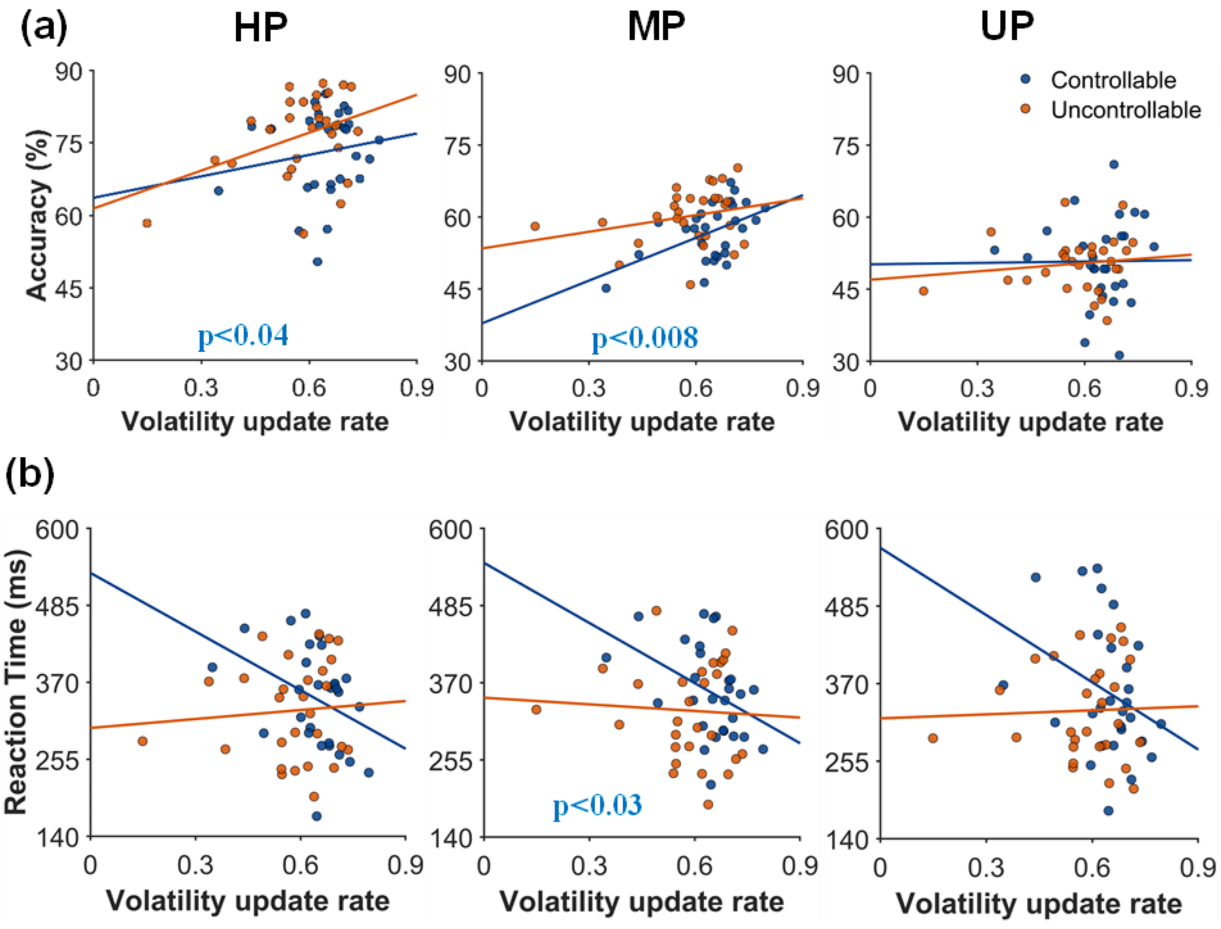
